# The Effects of Varying Gravito-inertial Stressors on Grip Strength and Hemodynamic Responses Across Gender

**DOI:** 10.1101/356154

**Authors:** Olivier White, Marie Barbiero, Nandu Goswami

**Affiliations:** INSERM UMR1093-CAPS, Université Bourgogne Franche-Comté, UFR des Sciences du Sport, F-21000, Dijon.; Chair of Physiology Unit, Otto Loewi Research Center for Vascular Biology, Immunology and Inflammation, Head of Research Unit: “Gravitational Physiology Aging and Medicine”, Neue Stiftingtalstrasse 6, D-5, Medical University of Graz, A 8036 Graz, Austria, EU

## Abstract

The body behaves as a global system with many interconnected subsystems. While the effects of a gravitational change on body responses have been extensively studied in isolation, we are not aware of any study that examined two types of body responses concurrently. Here, we examined how the neurocognitive and cardiovascular systems interact in this singular context and whether these combined responses are influenced by gender. Ten women and nine men underwent three 5-minute centrifugation sessions (2.4g at the feet, 1.5g at the heart) in which participants rhythmically moved a hand-held object for 20 seconds. Grip force and hemodynamic responses were continuously measured during centrifugation and rest periods. Our results show that men optimize the modulation between grip force and the destabilizing load force, but not women. Exposure to artificial gravity induced higher heart rate and mean arterial pressure in both genders compared to baseline. However, during exposure, only women decreased heart rate across sessions. Interestingly, we found that Finishers (N=13, mostly men) and Non-Finishers (N=6, mostly women) exhibited divergent patterns of hemodynamic responses. We also suggest that the lack of grip force adaptation reported in women can be linked to challenged hemodynamic responses in that population. Finally, by deriving a simple model to predict failure to complete the experiment, we found that mean arterial pressure was the most relevant dimension, and not gender. As artificial gravity is being proposed as a countermeasure in long-term manned missions, our results are particularly important but also deserve follow-up studies.

## Introduction

Orthostatic intolerance refers to the inability of a person to maintain upright posture without syncope, a transient loss of consciousness due to inadequate oxygen delivery to the brain (17). Orthostatic intolerance remains a problem upon return to Earth from the microgravity environment of spaceflight (4, 12). Almost every astronaut returning from space exhibits symptoms of cardiovascular deconditioning (3). However, there is a wide range of susceptibility to orthostatic intolerance after spaceflight, with some astronauts experiencing severe symptoms while others are minimally affected.

Artificial gravity (AG) administered during spaceflight or in ground-based analogues of spaceflight may prevent deconditioning in different physiological systems as well as prevent the development of orthostatic intolerance upon return to Earth. Current evidence from studies of ambulatory and deconditioned men and women indicate that 90 minutes of an individualized short- arm centrifuge AG profile significantly increased participants’ orthostatic tolerance limits via increased blood pressure and cerebral blood flow (3, 4, 8, 12–14). However, men and women respond very differently to these stressors (16). While this is still a topic of exploration, individualized AG seems to be a good countermeasure against orthostatic intolerance.

No other physiological system than the cardiovascular system responds more to changes in the gravitational vector within the body (12). On Earth, ambulatory subjects alternate between different gravitational stresses as they rise from sleep, sit, stand, walk and run before lying down to sleep again. In spaceflight, loss of these constant shifts in the body’s gravitational orientation leads to cardiovascular deadaptation that is evident in deterioration of reflexes that maintain blood pressure.

Beyond (cortical) blood flow alteration, AG also affects neurocognitive mechanisms at large (25). Maintaining operational motor and cognitive skills is also critical in these contexts. Importantly, the effects of AG on the body have always been considered in isolation. This view is however overly simplified since subsystems interact through constructive or destructive interferences. Using resting state fMRI and EEG, AG has been shown to alter cerebellar, sensorimotor and vestibular brain regions (6, 7, 18, 20, 21). More critically, gender-specific changes in cortical activation patterns during exposure to AG have also been reported using EEG (22). More precisely, that study revealed that alpha activity increased more in men than in women subject to the same AG. Usually, a decrease of alpha activity (and a parallel increase in beta activity) is associated with stress and arousal (5). This means that men and women react cognitively differently to AG as well and that this difference may impact the way each gender performs fine motor tasks.

Addressing research questions as to how AG affects the body holistically has recently been marked as a priority (25). Here, we investigated how cardiovascular responses interacted with the execution of a motor task in new challenging gravitoinertial contexts. The participants cyclically moved an instrumented object along the long body axis aligned with the gravitoinertial direction. We assessed how efficiently participants performed this simple manipulation task during rotation in a short arm human centrifuge. In addition, as most women participants could not complete the AG protocol, we also identified the profile of Non-Finishers in terms of cardiovascular and motor parameters.

## Materials and Methods

### Participants

Nineteen healthy men (n=9, 29.7±5.6 years old, 177.8±4.4cm, BMI 25.1±2.0kg/m^2^) and women (n=10, 27.6±4.6 years old, 165.1±4.8cm, BMI 21.9±1.9kg/m^2^) took part in this protocol. Each participant received a comprehensive medical examination from MEDES (French Institute for Space Medicine and Physiology), where the study was conducted, prior to participation. Inclusion criteria were: age between 20 and 40 years old, BMI<30kg/m^2^, normal clinical examination, normal electrocardiogram and arterial pressure. Exclusion criteria were the following: having taken part previously in hypergravity experiments in a short arm human centrifuge and having failed the medical exam. The experiment could be interrupted at any time upon participants’ request or his/her health status under constant monitoring by a medical supervisor. The experiment was interrupted during the last session (see *Experimental procedures)* for five women who showed signs of motion sickness and/or developed presyncope (Table 1). The signs for motion sickness included either performing the motor activity intermittently or verbally reporting so. Presynope was characterized by a sudden drop in systolic blood pressure to below 80 mmHg or a drop in heart rate by 15 beats per minuts (bpm) or the development of nausea, tunneling of vision or dizziness (elaborated in (11,15)).

**Table 1.** Participant details grouped by gender (F or M). The 3^rd^ column (N Sessions) provides the number of completed sessions. The 4^th^ column reports the total duration spent in the rotating centrifuge. In theory, the centrifuge rotated for 900 seconds (3×5 minutes, see Methods). The 5^th^ column indicates if physiological data (HR and MAP) could be analyzed. Finishers (reached the end of the experiment) and Non-Finishers (did not reach the end of the experiment or had to skip some conditions) are categorized in the last column.

| Participant | Gender | N Sessions | Exp. Duration (s) | Phys variables available | Finisher |
| --- | --- | --- | --- | --- | --- |
| P01 | F | 2 | 202 | no | No |
| P04 | F | 3 | 592 | yes | Yes |
| P06 | F | 3 | 857 | no | Yes |
| P10 | F | 3 | 829 | no | Yes |
| P11 | F | 1 | 260 | yes | No |
| P12 | F | 2 | 100 | yes | No |
| P13 | F | 3 | 605 | yes | Yes |
| P14 | F | 2 | 295 | yes | No |
| P18 | F | 1 | 83 | yes | No |
| P02 | M | 3 | 710 | yes | Yes |
| P03 | M | 3 | 767 | no | Yes |
| P05 | M | 3 | 525 | yes | Yes |
| P07 | M | 3 | 865 | no | Yes |
| P08 | M | 3 | 434 | yes | Yes |
| P09 | M | 2 | 501 | no | No |
| P15 | M | 3 | 645 | yes | Yes |
| P16 | M | 3 | 873 | yes | Yes |
| P17 | M | 3 | 342 | yes | Yes |
| P19 | M | 3 | 528 | yes | Yes |

The study was conducted in accordance with the ethical practices stipulated in the Declaration of Helsinki (1964). Ethics approval was obtained by MEDES (2014-A00212-45). All participants signed the informed consent form, which is stored at MEDES.

### Experimental procedures

Each participant was placed in a supine position on the centrifuge (Fig. 1). The participant rested his/her head on a thin pillow with the feet supported by a rigid metallic platform. The participant was equipped with headphones in order to maintain contact with the operator in the control room. An opaque ventilated box above the head prevented visual feedback of the environment.

**Figure 1.**
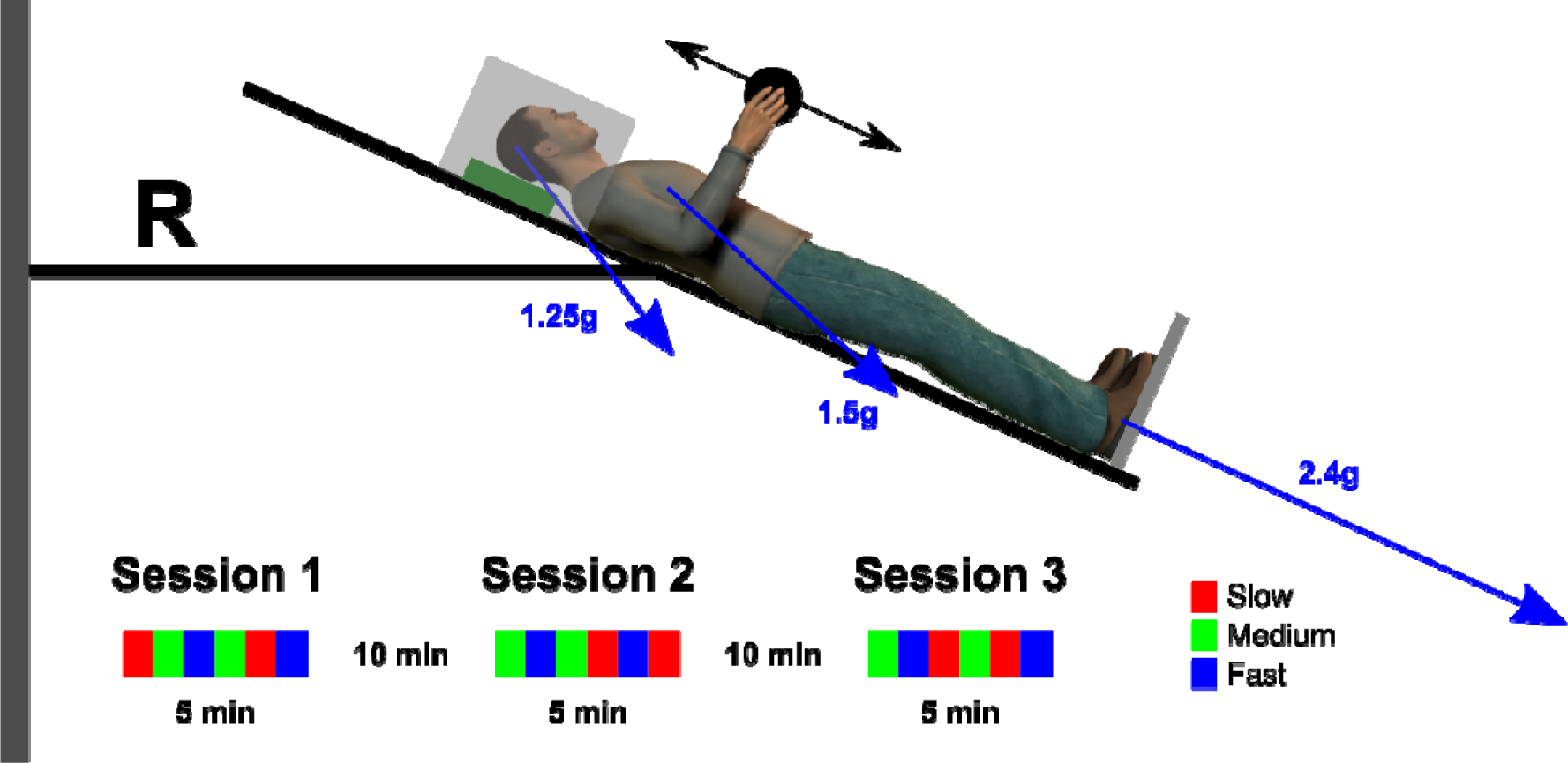
Scaled sketch of the participant in the centrifuge. The leftward grey vertical rectangle represents the axis of rotation of the centrifuge (angular velocity of —) The bed was tilted by −24 degrees and positioned such that the elbow joint was at distance R=1.63m from the axis of rotation. The participant was supine on the bed, her/his head resting on a cushion (green rectangle) and the feet supported by a metallic plate (grey line). Constant contact between the participant and the operator was maintained through headphones. The nacelle has local dark environments to eliminate the influence of external visual stimuli (transparent light grey rectangle). Gravitoinertial vectors depend both on Earth gravity and centripetal acceleration and vary in magnitude and direction (blue arrows, Head: 1.25g, −53deg; Heart: 1.5g, −42deg; Feet: 2.4g, −24.6deg). The two-headed arrow represents the trajectory of the object (black disk) in the sagittal plane. The lower inset illustrates a complete experimental schedule composed by three sessions. Each color corresponds to a different movement pace condition (see legend).

### Protocol

Participants underwent three centrifugation sessions each lasting for five minutes with the head tilted to 24 degrees upward (Fig. 1). This configuration was adopted to match technical requirements in a complementary study to maximize g-gradients along a short movement amplitude parallel to main body axis (2). Participants received 2.4g at the feet, which approximates to 1.5g at the heart and 1.3g at the head. Figure 1 describes the mechanical configuration and the gravitoinertial vectors. Each session was separated by 10-minute breaks during which the centrifuge nacelle was brought back to supine position and each participant rested quietly in the horizontal position.

### Object manipulation task

During centrifugation, and following a signal from the operator, each participant performed rhythmic upper arm movements in the sagittal plane with an instrumented object held in precision grip. The device recorded the 3d forces applied by the index and thumb finger as well as 3d accelerations of the object (detailed in (2)). Movement pace was provided by a metronome controlled by the operator that emitted 2 auditory signals per cycle, one at the top and one at the bottom of the trajectory. The signal was routed via headphones to the participant’s ears. Three paces *(Slow=*0.7Hz, *Medium=*1Hz and *Fast=13Hz)* were presented twice each for 30 seconds (3 paces x 2 repetitions x 30 seconds = 3 minutes). Pauses of about 20 seconds separated movement conditions in order to prevent fatigue. Pace order was randomized and counterbalanced across participants. The design ensured that participants were comparable across sessions in terms of total mechanical work performed. At the end of each session, the nacelle went back to idle position. All signals were continuously sampled at 200Hz and stored on a computer laptop strapped on the centrifuge.

### Hemodynamic recording

Continuous heart rate and arterial pressure were monitored using non-invasive photoplethysmography (Portapres: FMS, the Netherlands). The Portapres finger cuff was placed on the resting hand during the task. During tilting, it was ensured that finger measuring the blood pressure was held at the level of the heart using a Velcro strap.

### Data collection

Since all participants did not complete the experiment, we recorded the individual total exposure to AG. In addition, participants were categorized according to whether they could complete at least a few movement cycles in each condition and each session (Finisher) or not (Non-finisher). All Non-Finisher completed 1 or 2 sessions.

### Data processing and analysis

We continuously recorded 3d acceleration of the object (mass, m) and the grip force applied by the index finger and the thumb on the object (force orthogonal to the object’s surface). Movements induced inertial load forces (LF) of different magnitudes depending on oscillation frequency, or, object acceleration (a) according to *LF =* mg + ma, where g is gravitational acceleration (g=9.81 ms^−2^). We calculated mean grip force during each oscillation phase. Force signals were smoothed with a zero phase-lag autoregressive filter (cutoff 10Hz). A trial was defined as a series of cyclic movements. On average, per trial, participants performed 19.5 cycles for 0.7Hz (SD=6.9), 20.9 cycles for 1Hz (SD=2.2) and 26.3 cycles for 1.3Hz (SD=2.5).

For each session, we stored average heart rate (HR) and MAP values for 10-second epochs at baseline (before the centrifuge started), early after the centrifuge started (once HR was stabilized) and late (before the centrifuge ramped down to idle position). Furthermore, to quantify how HR and MAP varied over time, we fitted a linear function through these time series and analyzed the slopes of the regression lines.

### Statistical analysis

Matlab (The Mathworks, Chicago, IL) was used for data processing and statistical analyses. The force and physiological variables were averaged per participant and condition. Data were normally distributed as tested by the Shapiro-Wilk test (force: all p>0.632; physiological: p>0.05). We grouped the data according to gender (men and women).

We used a Fast Fourier Transform to extract the main frequency component of the acceleration profile for each trial. For each group, we used a two-way repeated measures ANOVA with a within-participant factor *session* (1^st^, 2^nd^ or 3^rd^) and *frequency* (0.7Hz, 1Hz, 1.3Hz). Post hoc tests were performed with Scheffé tests. Differences between groups were assessed with independent two-tailed t-tests. We report partial eta-squared values for significant results (p<0.05, corrected for multiple comparisons with Greenhouse-Geisser) to provide indication on effect sizes.

## Results

Participants cyclically moved an instrumented object while exposed to AG. We measured hemodynamic responses (through heart rate and arterial pressure) and how efficiently participants performed this motor task (through grip force). The aim of our investigation is to assess whether a fine motor action interacts with cardiovascular responses and how these responses are affected by gender in challenging stimulations.

### Gender differences in the dynamics of prehension

Participants adopted a pace that approached the instructions. Independent t-tests revealed no difference between actual and theoretical paces of 1Hz (t_24_=1.6, p=0.133) but faster paces than instructed in the slowest condition (t_24_=2.3, p=0.034, 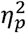) and slower paces than prescribed in the fastest condition (t_24_=2.1, p=0.045, 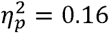). The ANOVA revealed significant effects of instructed frequency on pace (F_2,22_=69.7, p<0.001, 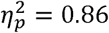) and mean grip force (F_2_,_22_=11.4, p<0.001, 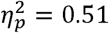).

Men exerted 40.7% larger grip forces than women overall (t_17_=2.5, p=0.024, 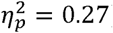). The efficiency of grip force adjustments is usually measured by computing the correlation between load force and grip force (9, 10, 23). This process is illustrated in Figure 2 A-B. The slope of the linear fit between load and grip forces quantifies the grip force increment (y axis) for a given load force increment (x axis) and the offset captures a safety margin, i.e. the force that would have been applied when LF=0 (see Fig. 2A-B). Men had significantly larger offsets than women (Fig. 2D, t_17_=2.3, p=0.036, 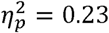) but gains and the quality of the fit, good on average (r=0.63), were similar (Fig. 2C, gain: t_17_=0.4, p=0.694 and Fig. 2E, correlation: t_17_=1.2, p=0.237). When each group was analyzed separately, the ANOVA only reported a significant effect of frequency in men on the offset (F_2,6_=10.3, p=0.031, 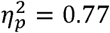) and on correlation (F_2,12_=10.6, p=0.003, 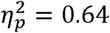). In both cases, a Scheffé post hoc shows that these variables increase with frequency but only in sessions 1 and 2 (offset: p<0.038; correlation: p<0.017). To sum up, our data demonstrate that the grip synergy slightly improves in men but not in women.

**Figure 2.**
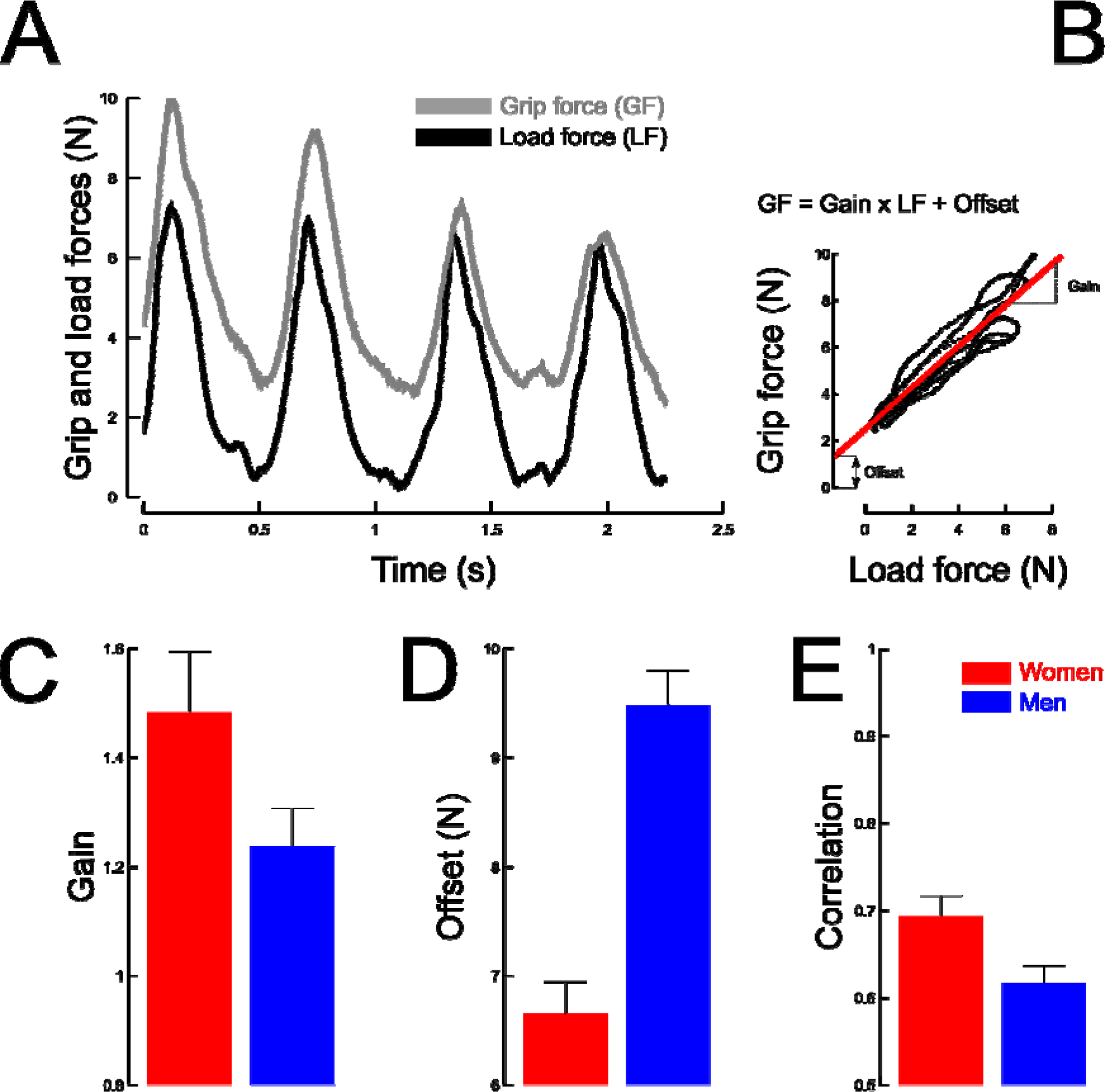
Main variables quantifying the grip force (GF) and load force (LF) synergy. (A-B): Grip force (gray line) parallels load force (black line) when a mass is moved cyclically along the vertical gravitational axis. (A) Depicts four cycles of movement over time. (B) Highlights the very good linear correlation between these GF and LF as a solid red line. The “Gain” and the “Offset” quantify this correlation. (C-E) Average values of gains (C), offset (D) and reliability of the linear relationship (E) plotted separately in women (red) and men (blue). Error bars represent SEM.

### Gender differences in heart rate and mean arterial pressure responses

Figure 3 depicts heart rate (HR, upper row) and mean arterial pressure (MAP, lower row) across sessions (1, 2 or 3) and separately in men (blue bars) and women participants (red bars). These variables are plotted separately at baseline (left column) and during the exposure to AG (right column). All participants increased their HR and MAP between baseline and AG exposure (HR: compare upper panels in Fig. 3, t_12_=20.1, p<0.001, 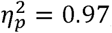; MAP: compare lower panels in Fig. 3, t_12_=11.6, p<0.001, 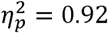). At baseline, the ANOVA did not show any significant effect of session on HR (Men: F_2,8_=0.4; p=0.653; Women: F_2,2_=7, p=0.229) or MAP (Men: F_2,8_=0.2; p=0.657; Women: F_2,2_=0.6; p=0.594). However, during AG exposure, women participants decreased their HR across sessions (F_2,2_=34.6, p=0.028, 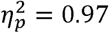) while men participants remained stable (F_2,8_=1.1, p=0.384). No effect of session was found in MAP during AG exposure across genders (all F<1.8, all p>0.507).

**Figure 3.**
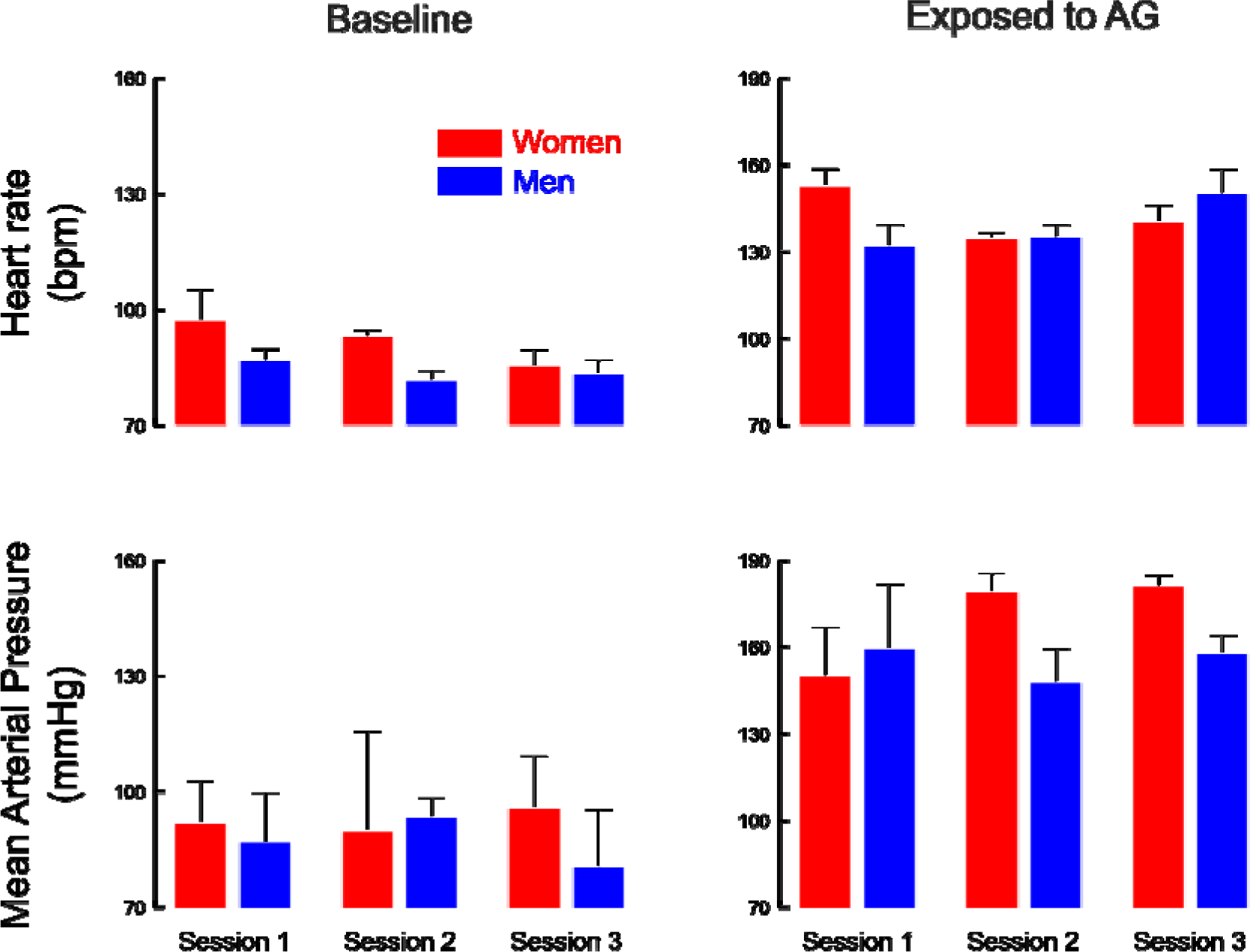
Heart rate (top row) and mean arterial pressure (lower row) at baseline (left column) and during exposure to AG (right column). Data are presented separately in men (blue bars) and women participants (red bars) and across sessions (x-axis). Note that for the sake of comparability, Y scales are identical for heart rate and mean arterial pressure between baseline and exposed to AG conditions. Error bars represent SEM.

Each session lasted five minutes. We observed that the slope of HR (Fig. 4A) was not affected by *gender* (t_11_=0.787, p=0.448) or *sessions* (all F<2.4, p>0.296). However, slopes were all positive, meaning that HR increased during AG exposure (t_24_=6.1, p<0.001, 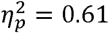). Similarly, the ANOVA did not report a significant main effect of *gender* (t_11_=0.991, p=0.343) or *session* on MAP slopes (Fig. 4B, all F<1.1, p>0.489). However, Figure 4B highlights very different trends of MAP slopes between men and women participants across sessions. Indeed, across sessions, MAP slopes increased in women and decreased in men. Independent t-tests that compare the MAP slopes within each session highlighted significant effects in *session 1* (t_10_=2, p=0.048, 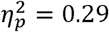) and *session 3* (t_8_=2.4, p=0.046, 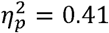) but not in *session 2* (t_7_=0.6, p=0.565).

**Figure 4.**
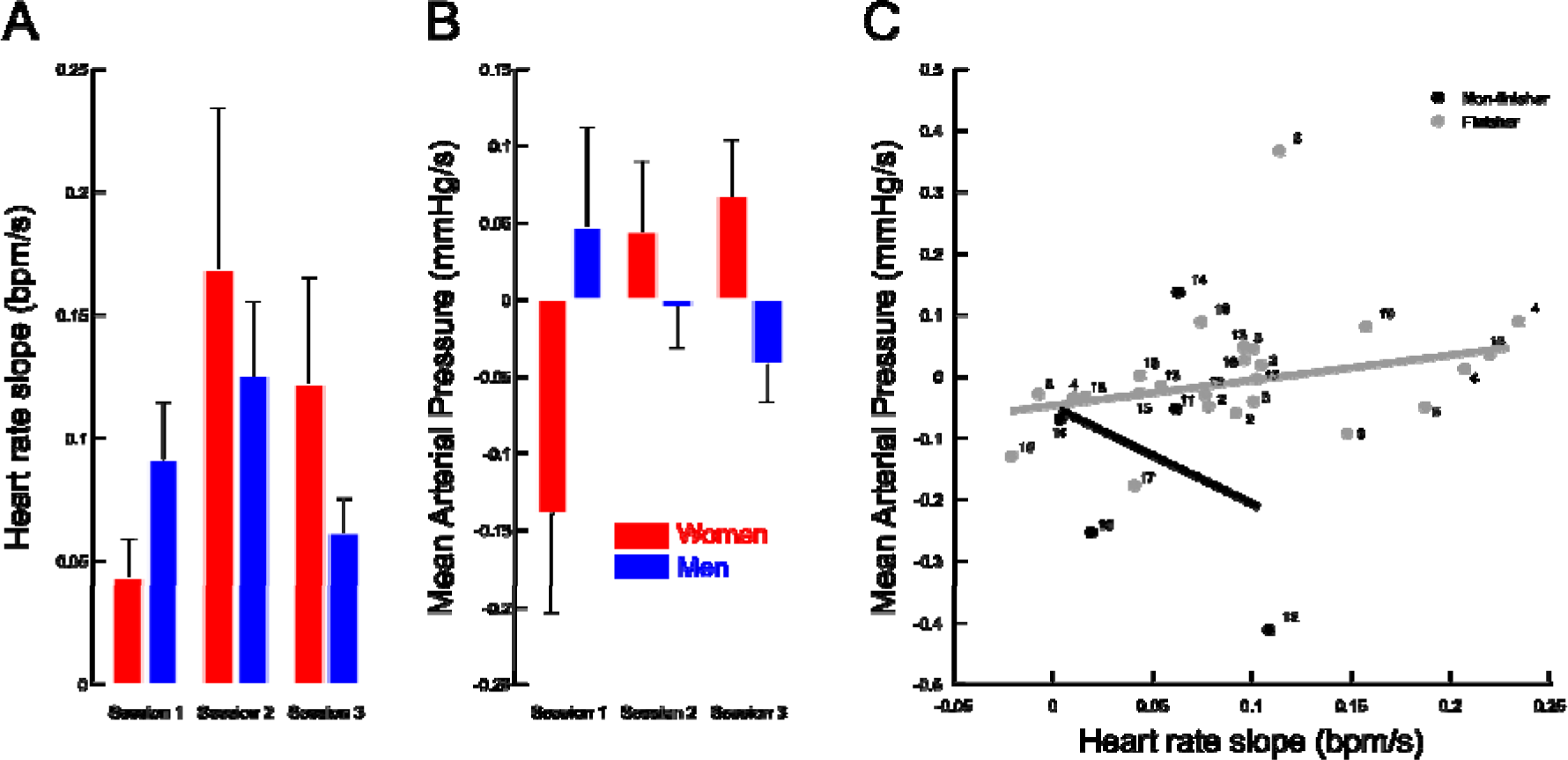
Variation of heart rate (A) and mean arterial pressure (B) over time, quantified by the slope of the linear regression between these variables and time. Men and women are separated and slopes are presented for each session. (C) Correlation between HR slope (x-axis) and MAP slope (y-axis) in the Non-Finishers (black disks) and in the Finishers (grey disks). Labels correspond to participant number.

Finishers (N=13, 69% men) and Non-Finishers (N=6, 83% women) depict contrasting patterns of HR and MAP. To quantify this relationship, we calculated the HR and MAP relative to baseline and correlated these values separately in Finishers and Non-Finishers. Figure 4C illustrates a significant correlation in Finishers (r=0.47, p=0.021) but not in Non-Finishers (r=-0.32, p>0.05). Altogether, this analysis shows that HR parallels MAP in men (mostly Finishers) while HR increases and MAP decreases in women (mostly Non-Finishers).

## Discussion

Our investigation aimed at identifying interactions between te cardiovascular and neurocognitive systems across genders while participants were exposed to AG. Participants performed oscillatory movements with a hand-held load at different paces. We observed that grip force was well adjusted overall and was optimized over sessions. Men exhibited smaller safety margins than women. During AG exposure, unlike men, women decreased their heart rate (HR) across sessions. There was no such effect found in mean arterial pressure (MAP). Finally, HR and MAP were increased in parallel in Finishers but showed opposite patterns in Non-Finishers, therefore yielding an index contrasting these two groups.

The slope of the linear fit between load and grip forces quantifies the grip force increment for a given load force increment (Fig. 2A-B). The larger the slope, the more efficient the grip adjustment. The offset of the fit reflects a safety margin that includes environmental factors and anxiety. For instance, this margin is slightly larger for a cup of coffee held stationary in an aircraft subject to unpredictable turbulences compared to the same object held in a quiet office. Overall, women exhibited significantly smaller offsets than men which reflects the spontaneous larger forces deployed in men. Both genders had larger safety margins than in familiar contexts which may be the sign of higher anxiety during AG exposure. Complex object manipulation tasks performed in new environements take time to adjust, which often initially translates in low gains, large offsets and poor correlations between grip and load forces (1). We observed significant effects of pace on the offset and on correlation but only in men. In both cases, these variables increased with frequency during sessions 1 and 2. This is a signature of an optimization of the motor task early in the experiment and for the two slowest frequencies. Note also that the medium frequency of 1Hz acted as an attractor for the slowest (uncomfortable) and fastest (too demanding) paces. Indeed, participants could not exactly match the prescribed frequency. Previous work showed that the gravito-inertial context strongly modulates natural movement frequency to optimize energy (24). Collectively, our data demonstrate that the grip synergy is roughly preserved in all conditions but significantly improves over sessions only in men.

### What determines whether a participant will be a finisher during an AG session?

Women had difficulties to complete the experiment. To our knowledge, we are not aware of any study that examined this aspect under the view angle of a motor task. We, therefore, investigated which parameters could be used to predict this result. As expected, Non-Finishers experienced AG a shorter amount of time than Finishers (t_17_=5, p<0.001, 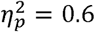). However, we only observed a trend when we split the data according to gender (t_17_=1.7, p=0.103), which means this criteria is not sufficient to state that sex is a predictor of being a finisher or not. Therefore, more parameters than gender should be included in the model. To investigate this further, we ran a principal component analysis separately in Finishers and Non-Finishers. We, therefore, included the following variables in the analysis: BMI, age, grip force offset, mean HR and mean MAP during AG exposure. We also included the product between slopes of HR and MAP in order to capture the divergent nature of these time series. As a result, Finishers required only two dimensions to explain 96% of the variance. The first dimension (76%) is positively dependent on MAP (coef=0.95) and the second dimension (20%) is a linear combination of HR (coef=0.85), grip force offset (coef=0.38) and age (coef=-0.34). In Non-Finishers, 93% of the variance was obtained using the first two dimensions as well. The first dimension (64%) is again positively dependent on MAP (coef=0.83) but now also negatively dependent on HR (coef=-0.54). The second dimension (29%) is a linear combination of HR (coef=0.81), MAP (coef=0.47) and grip force offset (coef=-0.31). To sum up, this holistic analysis shows that MAP is the most salient variable to predict failure to complete the experiment. Several studies have previously shown that blood pressure is a primarily related variable during stress application (19).

Furthermore, absolute heart rate seem also to be important when differentiating the two groups. Indeed, Finishers showed mean HR with a positive coefficient while Non-Finishers had a negative weight. Interestingly, the divergent nature of the HR and MAP time series could not allow classification of the two groups into Finishers and Non-Finishers. It appears that when categorizing participants into these groups, it is the absolute values of HR and MAP that are relevant rather than whether MAP and HR increase together or go in opposite directions over time.

Understanding mechanisms of neurocognitive changes and cardiovascular responses induced by AG application, an important countermeasure proposed for missions to e.g. Moon or Mars, is necessary. Furthermore, our results will provide insights into mechanisms which predisposes some persons to presyncope but not others during AG exposure. Finally, as the number of women crewmembers increases, the importance of studies designed to highlight major differences in cognition, handgrip strength and cardiovascular regulation in men and women during space missions also grows.

### Limitations

Our experimental context is challenging technically but also demanding for the participants. The data obtained in relation to cardiovascular measurements should be interpreted with caution as only two women with valid physiological data could finish the experiment. Similarly, the non-significant correlation seen in Figure 4C could be attributed to the small sample size. More studies are required to investigate this further.

## Acknowledgments

We wish to thank the MEDES team for their excellent efforts in helping us with the passing of the ethical agreement, recruitment of participants and for carrying out the experiment on the centrifuge. We are also grateful to the participants who took part in these experiments.

## Conflict of interest

The authors declare no conflict of interest.

## Funding support

This research was supported by the « Centre National d’Etudes Spatiales » grant 4800000665 (CNES), the « Institut National de la Santé et de la Recherche Médicale » (INSERM), the « Conseil Général de Bourgogne » (France) and the « Fonds Européen de Développement Régional » (FEDER).

